# UVC inactivation of pathogenic samples suitable for cryoEM analysis

**DOI:** 10.1101/2021.07.06.451241

**Authors:** JS Depelteau, L Renault, N Althof, CK Cassidy, LM Mendonça, GJ Jensen, GP Resch, A Briegel

**Affiliations:** Department of Microbial Sciences, Institute of Biology, Leiden University, Sylviusweg 72, 2333 BE Leiden, The Netherlands; Netherlands Center for Nanoscopy (NeCEN), Leiden University, The Netherlands; Department of Biochemistry, University of Oxford, Oxford, United Kingdom; Biology and Bioengineering Department, California Institute of Technology, Pasadena, CA, USA; Biology and Bioengineering Department, California Institute of Technology, Pasadena, CA, USA and Department of Chemistry and Biochemistry, Brigham Young University, Provo, UT USA; Nexperion e.U. – Solutions for Electron Microscopy, Vienna, Austria

## Abstract

Cryo-electron microscopy has become an essential tool to understand structure and function of biological samples, from individual proteins to whole cells. Especially for pathogens, such as disease-causing bacteria and viruses, insights gained by cryo-EM can aid in developing cures. However, due to the biosafety restrictions of human pathogens, samples are often treated by chemical fixation to render the pathogen inert, affecting the delicate ultrastructure of the sample. Alternatively, researchers use *in vitro* or *ex vivo* models, which are non-pathogenic but lack the complexity of the pathogen of interest. Here we show that ultraviolet-C (UVC) radiation at cryogenic temperatures can be used to eliminate or dramatically reduce the infectivity of two model organisms, a pathogenic bacterium (*Vibrio cholerae*) and a virus-like particle (the ICP1 bacteriophage). We show no discernable structural impact of this treatment of either sample using two cryo-EM methods: cryo-electron tomography (cryo-ET) followed by sub-tomogram averaging (STA), and single particle analysis (SPA). Additionally, we applied the UVC irradiation to the protein apoferritin (ApoF), which is a widely used test sample for high resolution SPA studies. The UVC-treated ApoF sample resulted in a 2.1 Å structure that did not reveal any discernable structural damage. Together, these results show that the UVC irradiation dose that effectively inactivates cryo-EM samples does not negatively impact their structure. This research demonstrates that UVC treatment is an effective and inexpensive addition to the cryo-EM sample preparation toolbox.

## Introduction

Cryogenic electron microscopy (cryo-EM) has emerged as a powerful technique for determining the structural characteristics of individual proteins, protein complexes, whole viruses, and even intact cells. The major advantage of this method is that it does not require any of the potentially artifact-inducing preparation steps needed for traditional transmission electron microscopy, such as dehydration, staining, or plastic embedding. Instead, the samples are simply flash-frozen into a glass-like ice (vitrified) and remain in a near-native state. Such samples can then be directly imaged using cryo-EM, providing high resolution structural information which in turn enables scientists to answer a wide range of specific biological questions.

The power of this technique is becoming increasingly apparent. For example, cryo-EM has given important insight into how components of SARS-CoV-2, the coronavirus causing the COVID-19 epidemic, interact with host cells, potential drugs, and candidate vaccines (for example, Juraszek et al., 2021; Ma et al., 2021; Subramaniam, 2020; Wolff et al., 2020; Yao et al., 2020). However, many of the structure-related studies resulted from either *in vitro* expressed proteins, isolated proteins from inactive virus, or a virus from the same family but lacks pathogenicity for humans. In other words, the information, while important to the understanding of how this virus infects its host and how to treat it, lacks structural information from the actual pathogen in a natural setting.

This is not only a limitation for the virus, but it is also the case for pathogens in general: the sample preparation for cryo-EM by flash-freezing is optimized to minimize structural damage. Therefore, precautions according to the specific biosafety level of any given sample need to be maintained throughout the entire sample preparation and imaging workflow. This limits the applicability of cryo-EM for obtaining structural information from such infectious pathogens, as cryo-EM facilities may not be equipped or approved to work with pathogens of biosafety level 2 or higher, or researchers are limited to specialized centers which host specialized, dedicated equipment such as a Titan Krios within a BSL2 or higher environment. This is a limiting factor for the research community’s ability to study these medically important organisms in real-life scenarios.

A recent study by Jin et al. (2015) demonstrated that ultraviolet (UV) irradiation of cryopreserved mouse embryonic fibroblasts has little-to-no effect on elemental distribution of the frozen hydrated samples. With similar ideas, we began testing whether UV irradiation could be applied to pathogens without compromising the ultrastructure of the samples. We tested this by applying ultraviolet-C (UVC) irradiation to inactivate two pathogenic organisms. UVC irradiation was chosen because of its well-described ability to inactivate pathogens (Boyce & Donskey, 2019; Welch et al., 2018). Inactivation occurs by altering the DNA of the organism, leading to the prevention of transcription and replication (Britt, 1996; Pattison & Davies, 2006). UVC irradiation is also known to impact proteins, mainly by disulfide bond breakage and the creation of reactive oxygen species (Durchschlag et al., 1996; Krisko & Radman, 2010). A combination of the above effects leads to the loss of viability of the organism, but the detailed effects on the structure of the pathogen are not well documented.

Here we describe a simple, inexpensive proof-of-principle prototype for administering UVC irradiation to cryo-EM samples under cryogenic conditions. We initially tested this device on two model organisms: the pathogenic bacterium *Vibrio cholerae* and the bacteriophage ICP1, a virus-like particle that infects *V. cholerae*. These organisms were chosen to test the applicability of the UVC inactivation for two different cryo-EM methods. The bacterium was used for cryo-electron tomography (cryo-ET) and sub-tomogram averaging (STA) workflows. These methods are used to study heterogeneous samples like whole bacterial cells and typically results in resolutions limited to macromolecular resolution (2-4nm, 20-40 Å). Single particle analysis (SPA) was used to evaluate the ICP1 bacteriophage sample, a method which allows the structure determination of identical particles to higher resolution (below 1nm, 10 Å). Finally, we used SPA of the protein apoferritin (ApoF) to determine structural changes at high (near-atomic) resolution that might result from UVC irradiation. ApoF is a common validation standard used in the EM community and resolutions of better than 3 Å are routinely achieved.

Here we show that UVC irradiation effectively inactivated both the pathogenic bacterium and the bacteriophage. We further show that the effects of this treatment on the structural information are non-discernable in the tested STA and SPA samples. Our study demonstrates that UVC irradiation of vitrified pathogen samples is a promising alternative to chemical fixation or the use of *in vitro* systems when access to a cryo-EM facility with sufficient biosafety clearance is not feasible. Combined with a simplified plunging device, it may be possible to freeze and UVC-treat samples locally, which can then be imaged at a lower safety level without compromising the structural information. In summary, this study provides significant evidence that UVC irradiation of cryo-EM samples may be a viable solution for structural investigations of a broad range of pathogen with cryo-EM.

## Methods and materials

### Bacteria and bacteriophage strains and conditions

*Vibrio cholerae* strains A1552 wildtype (WT; rifampicin resistance, 100μg/ml), N16961 WT (streptomycin resistance, 100μg/ml), and N16961-TndsRed (gentamycin resistance, 10μg/ml) were provided by Dr. Melanie Blokesch (Ecole Polytechnique Fédérale de Lausanne, Switzerland). *V. cholerae* were grown overnight in lysogeny broth (LB) containing antibiotic at 30°C with shaking at 180rpm. Equal volumes of an overnight culture and sterile 50% glycerol or 10% DMSO were mixed, frozen in liquid nitrogen, and stored at −80°C until use.

The ICP1 bacteriophage was provided by Andrew Camilli (Tufts University, Massachusetts, USA). The ICP1 phage was propagated using *V. cholerae* N16961 WT as described previously (Silva-Valenzuela & Camilli, 2019). Briefly, 10^6^ ICP1 phage was added to a 50ml culture (starting OD_600_ = 0.2) containing 5μM CaCl_2_. The culture was incubated for 4-6hrs at 30°C with shaking at 180rpm and the bacteria was pelleted by centrifugation (5000 g at 4°C for 30 min) and discarded. The supernatant is filtered through a 0.20μm filter, subsequently ICP1 is precipitated out using 20% PEG8000/2.5M NaCl solution overnight. Finally, ICP1 is pelleted by centrifugation (3000 g at 4°C for 1hr) and dissolved in phage storage buffer (100mM NaCl, 10mM MgSO_4_, 10mM Tris-HCl (pH 7.5), 1mM EDTA) and stored at 4°C until use.

### Sample preparation

*V. cholerae* was streaked onto a selective LB plate and grown overnight at 30°C and stored at room temperature. The night before the freezing session, several colonies were resuspended in liquid LB containing antibiotic and grown overnight at 30°C with shaking at 180rpm. An aliquot of this culture was used for freezing as described below. For ICP1, an aliquot from the phage stock was used directly for sample freezing. To minimize lipid vesicles in the ICP1 sample, Tween-80 (Sigma Aldrich) was added to the UVC-treated sample shortly before freezing to a final concentration of 0.05%.

The bacterial and phage samples were prepared using a Leica EM GP (Leica Microsystems, Wetzlar, Germany). For the bacterial sample, 15nm gold beads was added to the bacteria prior to freezing (Cell Microscopy Core, Utrecht University, Utrecht, The Netherlands). For both samples, 3μl was applied to a glow discharged Quantifoil R2/2, 200 mesh Cu grid (Quantifoil Micro Tools GmbH, Jena, Germany), and incubated prior to blotting (30s for *Vibrio cholerae* or 10s for ICP1) at 18°C with approximately 90% relative humidity. The grid was blotted for 1s and automatically plunged into liquid ethane. Vitrified samples were transferred to storage boxes and stored in liquid nitrogen until use.

Human apoferritin (ApoF) sample was used at a concentration of 4 mg/mL and applied to glow discharged Quantifoil R2/2, 200 mesh grid before being double-side blotted for 3 seconds in a Vitrobot Mark IV (Thermo Fisher Scientific) and plunge frozen into liquid ethane. Frozen samples were stored in liquid nitrogen until UVC treatment and imaging.

### UV irradiation device and protocol

The UV irradiation device was constructed using easy-to-acquire materials. Working with our local fine mechanical department, we created a sandwich of three quartz glass slides (Figure S1A). Quartz glass was chosen because it does not interfere with UVC radiation, as shown with spectroscopy. This sandwich design allows for the accurate placement of the frozen grids during the UVC irradiation process (Figure S1B). A holder for the grid box and quartz slide sandwich was designed using styrofoam, and inserted tightly into a modified blue foam liquid nitrogen dewar (Figure S1C; Spearlabs Cryogenic Products). The styrofoam holder contains a shallow well sized to the assembled quartz glass sandwich, and is lined with aluminum foil to encourage omnidirectional irradiation (Figure S1C arrow). This assembly allowed for the loading and unloading of grids under liquid nitrogen conditions (Figure S1D), and it permitted the placement of the UVC light source directly above the samples.

The UVC light source (6W Germicidal Light T5 Tube UVC Sterilizer, Lcamaw via Aliexpress.nl) was mounted to the underside of a box to allow the placement of the blue sample assembly and a UVC sensor (UV Light Meter LS126C, Linshang Technologies, China) directly below the light source (Figure S1E). The UVC sensor was placed adjacent to the sample assembly at sample height to confirm irradiation and dose rate. At the sample, the dose rate from the light source reached a maximum of 490 μW/cm^2^, as determined with the final reading before discontinuing UVC treatment. Based on this setup, a box was designed to properly contain the light source, sensor, and sample assembly (Figure S1F).

Samples were irradiated as follows: During each irradiation experiment, four sample-containing grids were processed. Three of the grids are transferred to a position of the quartz grid holder. The fourth grid was left in the grid box with the lid closed to act as a control (grid box lid is opaque and grid remains perpendicular to the light source, preventing the passage of UVC to the sample). The liquid nitrogen level was then raised to a level of 0.5 cm above the sample, which we have marked on inside of the blue foam container (Jin et al., 2015). Prior to irradiation, the UVC light was switched on for one minute (prewarming step) and then switched off. Immediately after sample assembly and pre-warming, the lamp is placed over sample assembly, and the lamp switched on for the treatment time. For the bacterial sample, grids were treated on the carbon side only; for the bacteriophage sample, irradiation occurred on both the carbon and copper side of the grids (by inverting the sandwich in the liquid nitrogen/vapor layer). Once treatment was complete, the grids are transferred back to the grid box and stored in liquid nitrogen until the confirmation of inactivation and or imaging.

### CFU and PFU

To determine the exposure time for UVC treatment of *V. cholerae* or the ICP1 bacteriophage, the viability of the samples was determined by either colony forming units (CFU) for the *V. cholerae* WT, or plaque forming units (PFU) for the bacteriophage exposed to *V. cholerae* N16961-TndsRed. The treated grids were transferred to LB, pulse vortexed several times to resuspend the bacteria/phage and then subjected to serial dilution. CFUs were determined by plating 100ul of the serial dilution onto non-selective LB plates and incubated overnight at 30°C. The PFU were determined as follows: *V. cholerae* N16961-TndsRed containing soft agar was overlayed on square LB plates containing 10μg/ml gentamycin. 5μl from each dilution step was spotted onto the top layer and the plate left to grow overnight at room temperature. Clearance zones were noted the following day, and PFU were roughly determined. The duration of UVC exposure to inactivate greater than 99.99% of the phage was determined based on the amount of time needed to prevent ICP1 predation of *V. cholerae*, demonstrated by the lack of clearance zone.

### Imaging conditions

#### Cryogenic electron tomography

The grids containing vitrified, UVC-treated *V. cholerae* were clipped and loaded into a CS-corrected Titan Krios (Thermo Fisher Scientific (TFS)) equipped with a K2 direct electron detector and a post-column energy filter (Gatan, Inc) set to zero loss imaging with a 20eV slit. Targets were chosen based on the presence of the flagellar pole in a hole. This extracellular feature is a good indicator for the presence of the F6 chemotaxis array that is located at the same cell pole. A tilt series of each target was collected using SerialEM set to a dose symmetric tilt scheme between −54° and 54°, with 2° increments, with a pixel size of 3.49 Å (Hagen et al., 2017; Mastronarde, 2005). A defocus of −8μm and a cumulative dose of 140e-/Å were used as targets.

For the untreated data set, we used data collected during a previous session. This data was collected on a Titan Krios (TFS) microscope equipped with a K3 BioQuantum direct electron detector and energy filter (Gatan, Inc) set to zero loss imaging with a 20eV slit. Whole cells in a hole were selected as targets. Tilt series were collected using a bidirectional scheme between −60° and 60° with 2° increments with a pixel size of 5.86 Å. The target defocus was set to −8μm and the estimated total dose 170e-/Å.

#### Single Particle

Treated and untreated ICP1-containing grids were clipped and loaded into a Titan Krios (TFS) microscope equipped with a K3 BioQuantum direct electron detector and energy filter (Gatan, Inc) set to 20eV. Micrographs were collected using the TFS EPU software equipped with AFIS (aberration free image shift) with a pixel size of 0.685 Å in super-resolution mode. Defocus was set cycled between −1 and −3 μm, and a dose of 34e/Å^2^ per image.

Apoferritin samples were treated as described in Diebolder et al (Diebolder et al., 2021). Briefly, grids were loaded into a Titan Krios electron microscope (TFS) operated at 300kV, equipped with a Gatan K3 BioQuantum direct electron detector (Gatan, Inc). Movies with 50 frames and an accumulated dose of 50 electrons/Å2 were acquired in super resolution counting mode using EPU (TFS) at the magnification of 130,000X, corresponding to a calibrated pixel size of 0.328Å/pixel with a defocus range of −0.5 to −2.5μm. A total of 1,040 movies were collected at The Netherlands Centre for Electron Nanoscopy (NeCEN). Detailed data acquisition parameters are summarized in Supplementary Table S1.

### Data processing

#### Vibrio cholerae

Motion correction and tomogram generation were done using the IMOD image processing suite (Kremer et al., 1996). In brief, once the tilt series were motion-corrected using the alignframes function, the batchruntomo function was used to generate the initial tomograms to determine if the structure of interest was present in the cell (final bin = 3 without CTF correction; Mastronarde & Held, 2017). Tomograms containing the feature of interest were further processed by refining the bead model, or using patch tracking if insufficient beads were present. Subsequently, the boundary model was improved, CTF correction was applied, and a back-projected, SIRT-like filtered tomogram was generated (Xiong et al., 2009). A final bin of 2 was used for the UVC treated data (pixel size = 6.98 Å) and a final bin of 1 for the untreated data.

The resulting tomograms were used for STA using the Dynamo imaging suite (Castaño-Díez et al., 2012, 2017). Subtomograms were manually picked from chemotaxis array top views. Two rounds of iteration steps were performed, using a template generated from particles picked from a single tomogram, and a mask that encompassed a single hexagon. All final maps were calculated from weighted back-projection SIRT-like filtered tomograms. The resolution of the final maps were calculated using ResMap (Kucukelbir et al., 2014).

#### ICP1

Reconstruction of the ICP1 capsid was done using Relion 3.1.2 (Scheres, 2012). The data was binned by 4 for the initial particle picking, 2D, and 3D refinements. GPU enabled MotionCorr was used to correct particle movement as part of the Relion processing suite (Zheng et al., 2017) and gCTF was used for CTF estimation (Zhang, 2016). Initially, 150 phage full and empty heads were manually picked for 2D classification to generate reference templates for autopicking. Following autopicking and extraction, 2D classification was performed using the T4 phage head, PDB 8661, filtered to 70 Å as a reference (Chen et al., 2017). 3D classification was performed with a subset of particles using the 2D classification full head as a template. The full head class was auto-refined using “I” symmetry. Post processing used a mask generated by Relion. Particles were re-extracted with a pixel size of 1.37, 3D autorefine and postprocessing was repeated yielding the final structure.

#### Apoferritin

RELION-3.1 beta software was used for ApoF image processing (Scheres, 2012; Zivanov et al., 2018). Collected movies were subjected to beam induced drift correction using MotionCor2, the contrast transfer function was estimated by CTFFIND-4.1.18 (Rohou & Grigorieff, 2015; Zheng et al., 2017). RELION Gaussian picker was used to automatically pick 251,350 particles. After 2 rounds of 2D classification, false positive and contaminating features were discarded resulting in a 66,920 particles dataset. Ab-initio model generation followed by 3D classification and 3D refinement yielded a 2.6 Å map. Corresponding particles were subjected to CTF refinement for optical and beam-tilt and aberration correction, as well as per-particle defocus, and per-micrograph astigmatism correction followed by Bayesian polishing (Zivanov et al., 2019, 2020). A second 3D refinement was then performed yielding a 2.1 Å map.

Map resolution were estimated at the 0.143 criterion of the phase-randomization-corrected FSC curve calculated between two independently refined half-maps multiplied by a soft-edged solvent mask. Final reconstructions were sharpened and locally filtered in RELION post-processing. Maps were displayed using UCSF ChimeraX (Goddard et al., 2018).

#### Molecular modeling and simulations

An initial homology model of the major capsid protein was constructed using the Gp122 sequence and trRosetta webserver (Barth et al., 2020; J. Yang et al., 2020). A hexameric complex was assembled into each map by rigid docking using UCSF Chimera v1.13 (Pettersen et al., 2004). A 5-ns molecular dynamics flexible fitting (MDFF) simulation was performed on each hexamer using NAMD v2.13 and the cascade-MDFF protocol with default parameters (Phillips et al., 2005; Singharoy et al., 2016). Fittings were carried out with an MDFF coupling constant of 0.1 and symmetry restraints applied on the backbone atoms of each monomer. Structural visualization and trajectory analysis were performed in VMD v1.9.4 (Humphrey et al., 1996).

## Results

### UV inactivation of *Vibrio cholerae*

As an initial test of the UV inactivation device, we determined the UVC exposure time necessary to inactivate the pathogen, *V. cholerae A1552 WT*. Our experiments showed a significant reduction in vitrified, viable cells after 20 s of UVC exposure, and complete inactivation with 30 s of UVC exposure. Based on this information, we chose 30 s as the timepoint for inactivation and subsequent imaging.

Following the UVC treatment of the *V. cholerae* cryo-EM sample, the grids were transferred to a Titan Krios electron microscope for data collection. To determine the effect of the UVC irradiation, we collected tilt series of the flagellar poles of individual cells, with the aim of imaging the F6 chemotaxis array, which are located at the same pole. Chemotaxis arrays are well-studied by ECT and STA, and thus provided a good model system for determining of macro-level damage to cells. Following data collection and subsequent processing, tomograms with top views of the F6 chemotaxis array were identified and further processed using STA (Figure 1A/C). For the control sample, and using 219 particles, we were able to achieve a resolution of 22.6 Å (Figure 1B), whereas with the UVC-treated sample and 170 particles, we were able to achieve a final resolution of 26.7 Å (Figure 1D).

**Figure 1:**
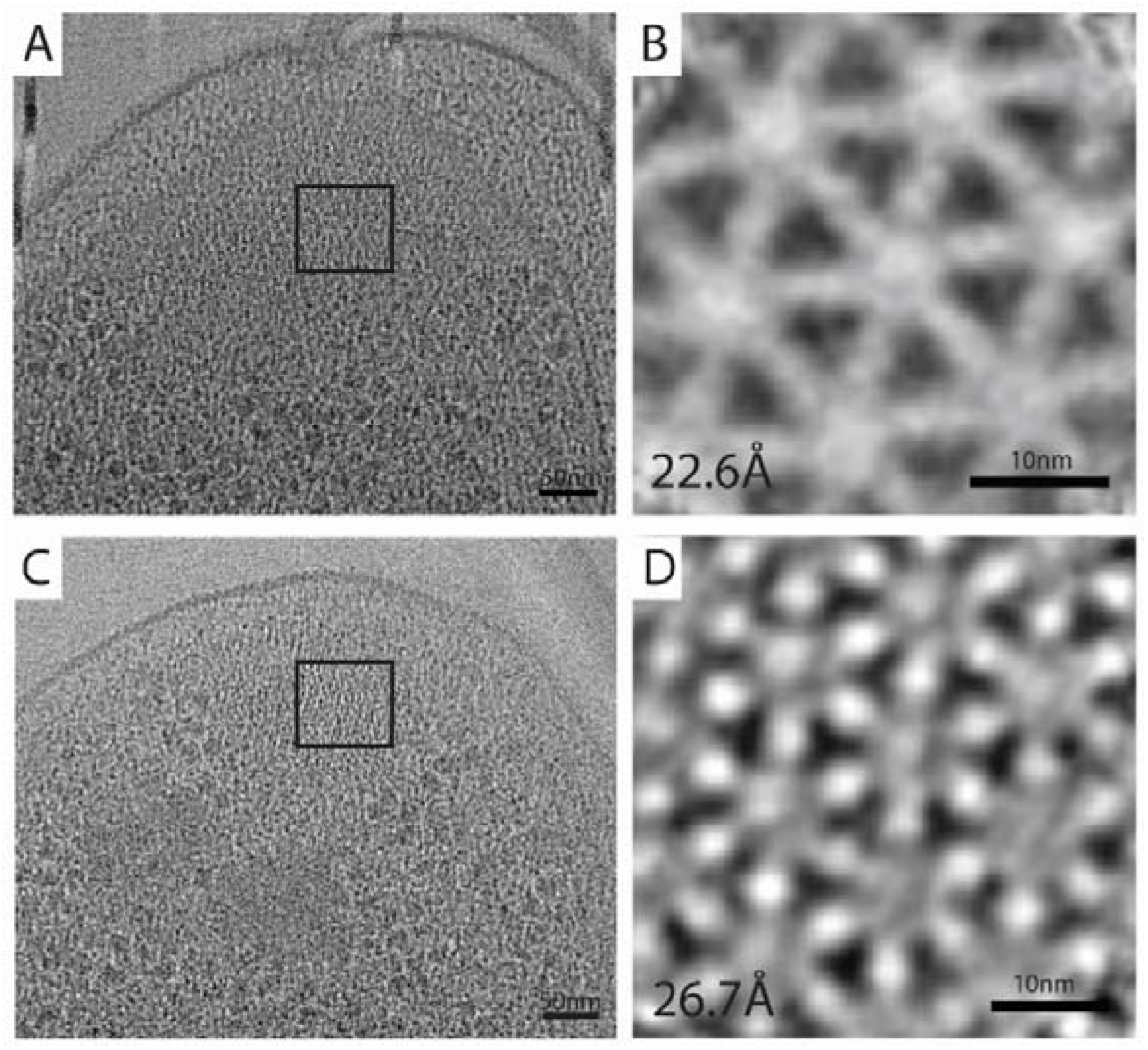
Effect of UVC irradiation on *Vibrio cholerae and its chemotaxis array*. Panels A and C show the flagellar pole region of untreated (A) and UVC-treated (C) *V. cholerae* cells, highlighting the top view of the F6 chemotaxis array (black box). Panels B and D are the results of subtomogram averaging of F6 chemotaxis cluster, showing the typical hexagonal arrangement of the trimer-of-dimer receptors in extended arrays.

### UV inactivation of ICP1 phage

The ICP1 bacteriophage was chosen as a model for a virus-like particle. We determined the UVC exposure time that was necessary to inactivate greater than 99.99% of the phage. This was determined to be 60 s for the carbon side and 30 s for the copper side (Figure 2A). We then UVC irradiated a new set of ICP1-containing grids, confirmed inactivation of phage, and used the remaining grids for imaging (Figure 2B).

**Figure 2:**
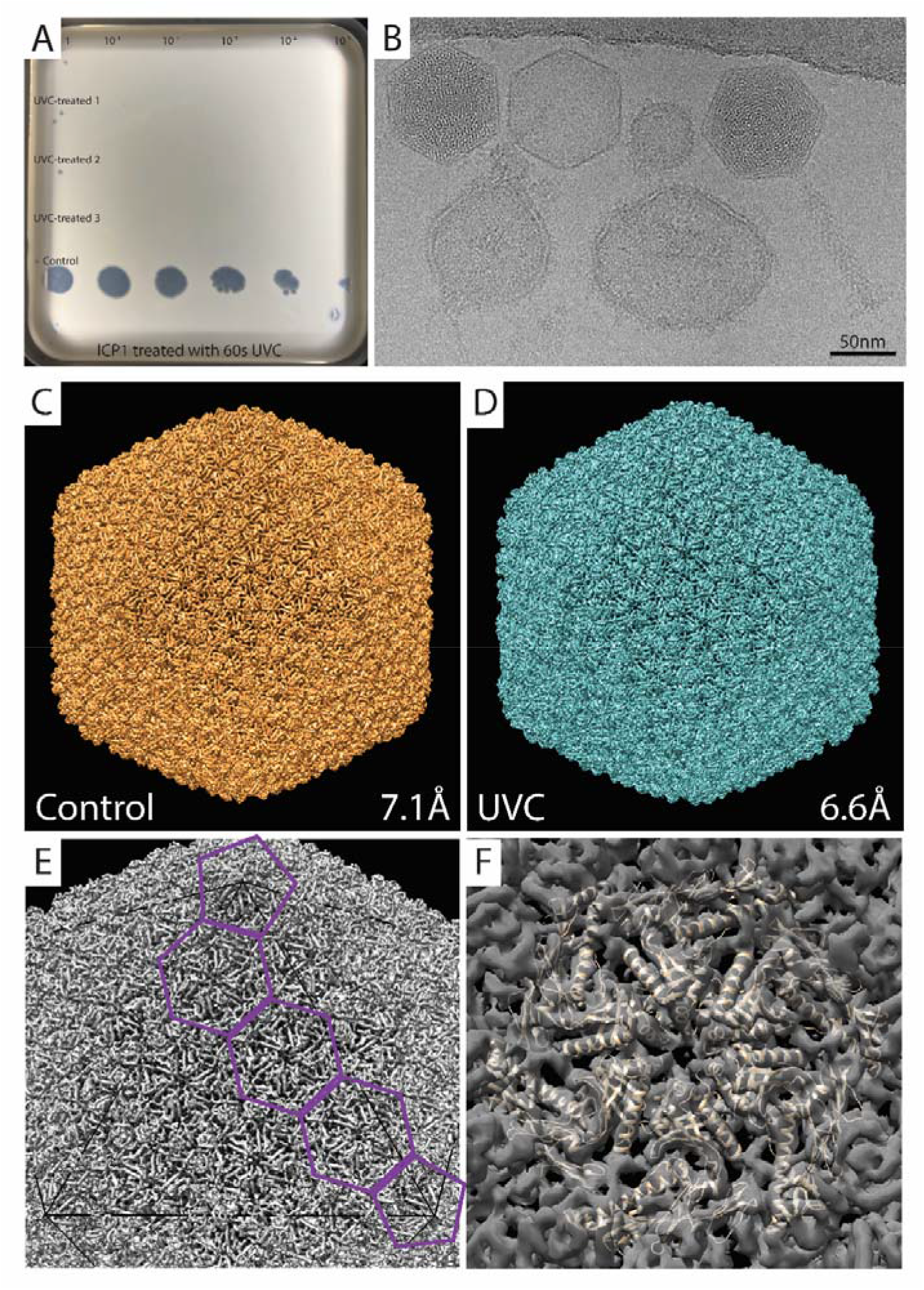
UVC treatment and SPA of ICP1. A. UVC treatment of ICP1 resulted in a significant reduction in viable phage. Rows 1-3 are UV treated and show 0-2 infective phages in three replicates when undiluted (dilution factor noted at the top of panel). Row 4 is a negative control, a grid containing ICP1 that remained in the grid box during treatment. B. A representative micrograph of UVC-treated ICP1 showing an assembled phage (far right), an empty and full head (middle/left), and liposomes (round, bottom). C & D. SPA analysis resulted in structures of the control and UVC treated ICP1 phage with a resolutions of 7.1 Å and 6.6 Å, respectively. E. Determination of the T number = 13, which a measure of the icosahedral symmetry. F. Flexible fitting of putative major capsid protein, ORF122, as a hexamer, and docked into the ICP1 capsid of the UVC-treated structure noted in D.

Following imaging, the number of particles and classes was determined before subsequent processing. For the UVC-treated sample, 6577 particles were separated into two classes: capsid containing DNA and empty capsids. This was done because previous research in our group had shown that the size of the empty capsid is slightly larger than the capsid containing DNA (unpublished results). Here, only capsids containing DNA were used for 3D classification and refinement. Ultimately, 2845 particles resulted in a final resolution of 6.6 Å. A similar methodology was used for a separate, untreated data set, which resulted in a final resolution of 7.1 Å using 4096 particles. A comparison of the two structures using Chimera:Fit in Map yielded a correlation value of 0.9815. The achieved resolutions allowed for the identification of the T number (T=13; Figure 2E; (Caspar & Klug, 1962)) which is a metric for the icosahedral symmetry, and the flexible fitting of the putative major capsid protein into the map (Figure 2F).

To compare the local quality of the two maps, we used each to structurally refine a hexameric assembly of major capsid proteins (Figure 2F). For this purpose, we carried out molecular dynamics flexible fitting (MDFF) simulations (Methods) (Singharoy et al., 2016), which use a density-derived potential to optimize model-map overlap. The refined hexamer models were nearly identical, possessing a backbone root-mean-square-deviation (RMSD) of 1.23 Å for all residues and 0.99 Å for non-loop residues. The backbone RMSD between the refined and initial hexamer models was 3.4 Å and 3.6 Å, respectively, for the untreated and UVC-treated maps. The structural information contained in both maps is therefore of comparable quality.

### UV inactivation of ApoF

Finally, we wanted to determine the effect of UVC irradiation at resolutions similar to x-ray crystallography, which allow the visualization of individual amino acids. To do this, we used the protein complex, apoferritin (ApoF). ApoF is commonly used in EM facilities to determine the performance of microscopes and it has been well characterized by the scientific community using cryo-EM and x-ray diffraction. Using the same UVC exposure time as ICP1, grids containing apoferritin were treated with UVC for 60s on the carbon side and 30s on the copper side. These grids were subsequently imaged and the data processed using SPA. Our experiments demonstrate that we were able to achieve a final resolution of 2.1 Å in the UVC-treated sample, which is on par with structures published in the Protein Data Bank (https://www.ebi.ac.uk/pdbe/node/1;.

A comparison of the UVC treated apoferritin with untreated samples and published structure found no difference between the two structures (Figure 3; EMD-3853 (unpublished), PDB5N27 (Ferraro et al., 2017)). A closer analysis of selected regions of the structure of the treated apoferritin sample highlight the structural preservation (Figure 3B,C).

**Figure 3:**
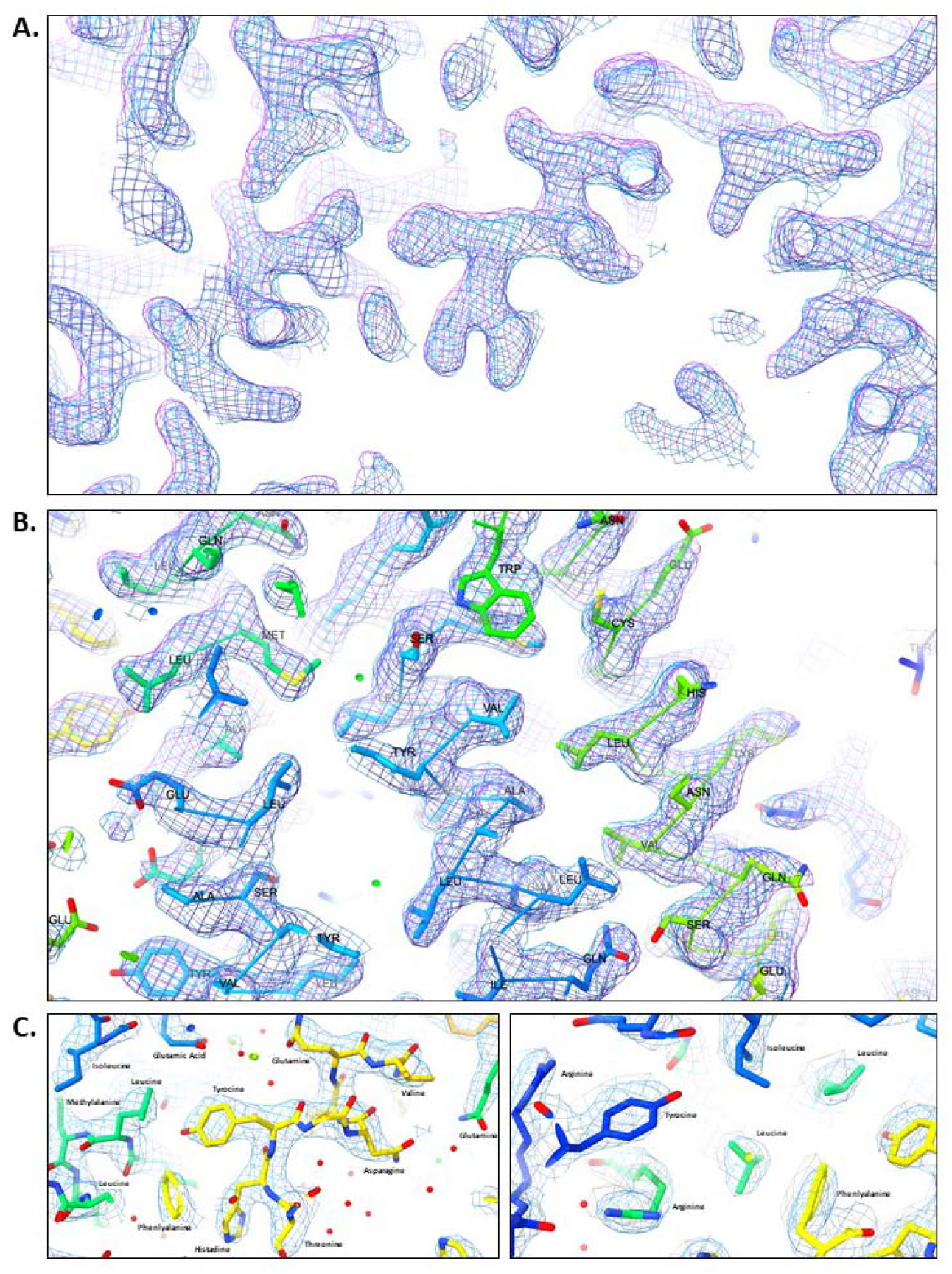
Representative ApoF amino acids show no difference between crystal structure and UVC-treated ApoF cryo-EM map. A. A comparison of the UV-treated ApoF map (blue) with a previously published, untreated structure (purple; EMD-3853) shows no difference between structures. B. The UV-treated ApoF map (blue) superimposed with a untreated map (EMD-3853) and fitted model (PDB-5n27) confirms no significant differences. C. Close-up view of UV-treated map with the fitted model (PDB-5n27), highlighting no difference between representative amino acids that might be impacted by irradiation.

## Discussion

UVC irradiation is a widespread technique used for disinfection (Buonanno et al., 2020; IES, 2020). In this paper, we demonstrate that UVC irradiation is also suitable for inactivating pathogens in cryogenic samples intended for structural analysis by cryo-EM. Our results confirm a significant reduction and or elimination of the ICP1 bacteriophage and its target organism, *V. cholerae*. We characterized the effects of this treatment by comparing cryo-EM data of control (untreated samples or previously published structures) with samples after treatment and inactivation. Here, we show at multiple scales that UVC irradiation has no discernable effect on the structural information of the biological cryo-EM samples.

Cryo-EM has become a valuable method for understanding bacterial cell biology. Using cryo-ET, the organization of bacterial cells and their molecular machines have become accessible for structural analysis. Thus, we began by determining the effects of UVC irradiation on a widely studied bacterium. We show that while the bacterial pathogen *V. cholerae* is completely inactivated by UVC, the cell’s ultrastructure is indistinguishable from untreated samples. More specifically, using the F6 chemotaxis array as an example, we show that the STA results (reaching resolutions between 2-3 nm) are similar to the published literature for complexes of similar size and in cells with similar thicknesses (Berger et al, 2021; Wolff et al, 2020; Yang et al, 2019; Muok et al, 2020).

We next wanted to test the suitability of this treatment method on a virus-like particle. We therefore tested the UVC treatment on ICP1, a bacteriophage that infects *V. cholerae*. Similar to the bacterial sample, the bacteriophage was also effectively inactivated using the UVC treatment of the cryogenic samples. The resolution reached here (^~^7 Å) allowed for the identification of individual alpha helices, turns within the protein, and groups of beta sheets. This information was used gain structural information about the ICP1 capsid, including its overall structure as demonstrated by its T number, a spike protein complex at the vertices (data not shown), and the docking of the putative major capsid protein into the capsid hexamer. This result further illustrated the suitability of the UVC inactivation method for low-resolution SPA of virus-like particles.

Finally, we demonstrated the applicability of this method for high-resolution SPA. At resolutions of better than 3 Å, structural damage would likely be visible by the lack of disulfide bonds and damage to aromatic amino acids. Our experiments show that UVC irradiation had no impact on our ability to obtain a final structure of ApoF at a resolution of 2.1 Å (Figure 3). This resolution is comparable to other reported structures (EMD-3853 (unpublished), PDB5N27 (Ferraro et al., 2017)). A closer examination of the treated and untreated data did not reveal any noticeable differences between the two structures (Figure 3).

This research did not come without some challenges. For instance, our initial experiments to determine PFU used the same strain of V. cholerae as used for the cryo-ET and STA. However, we found that this strain and antibiotic combination gave inconsistent PFU counts and thus necessitated a switch to a strain more similar to the ICP propagation protocol. When working with the ICP1 sample, we also found many factors could influence the reduction in infectious particles, including contamination from forceps, bending of the grids during transfer, and the presence of outer membrane vesicles/lipid containing particles in the sample preparations. Thus, it will be important to determine the needs for their specific sample prior to imaging.

Together, this research demonstrates that UVC irradiation of vitrified samples can provide significant structural information for a variety of samples and across scales. We believe this information is especially useful for labs who study pathogens that may not be approved for imaging at their local, regional, or national facilities. We demonstrate that UVC treatment of the samples directly on the grid preserves the structural information while rendering the pathogen with reduced or no infectivity. It is important to note that the inactivation protocol will have to be adapted to each pathogen of interest. Regardless, this inactivation method provides an affordable, straightforward method to inactivate pathogens for cryo-EM studies and will be useful for all laboratories lacking access to cryo-EM facilities that are certified for higher biosafety level microorganisms.

## Acknowledgments

We are grateful to Davi Ortega for helpful discussions. We also wish to thank Dr. Weng Yang and Dr. Willem Noteborn of NeCEN for assistance during data collection, and Gert Koning (Department of Fine Mechanical) for assistant with the construction of the prototype UVC inactivation box.

## Supplemental Figures

**Figure S1:**
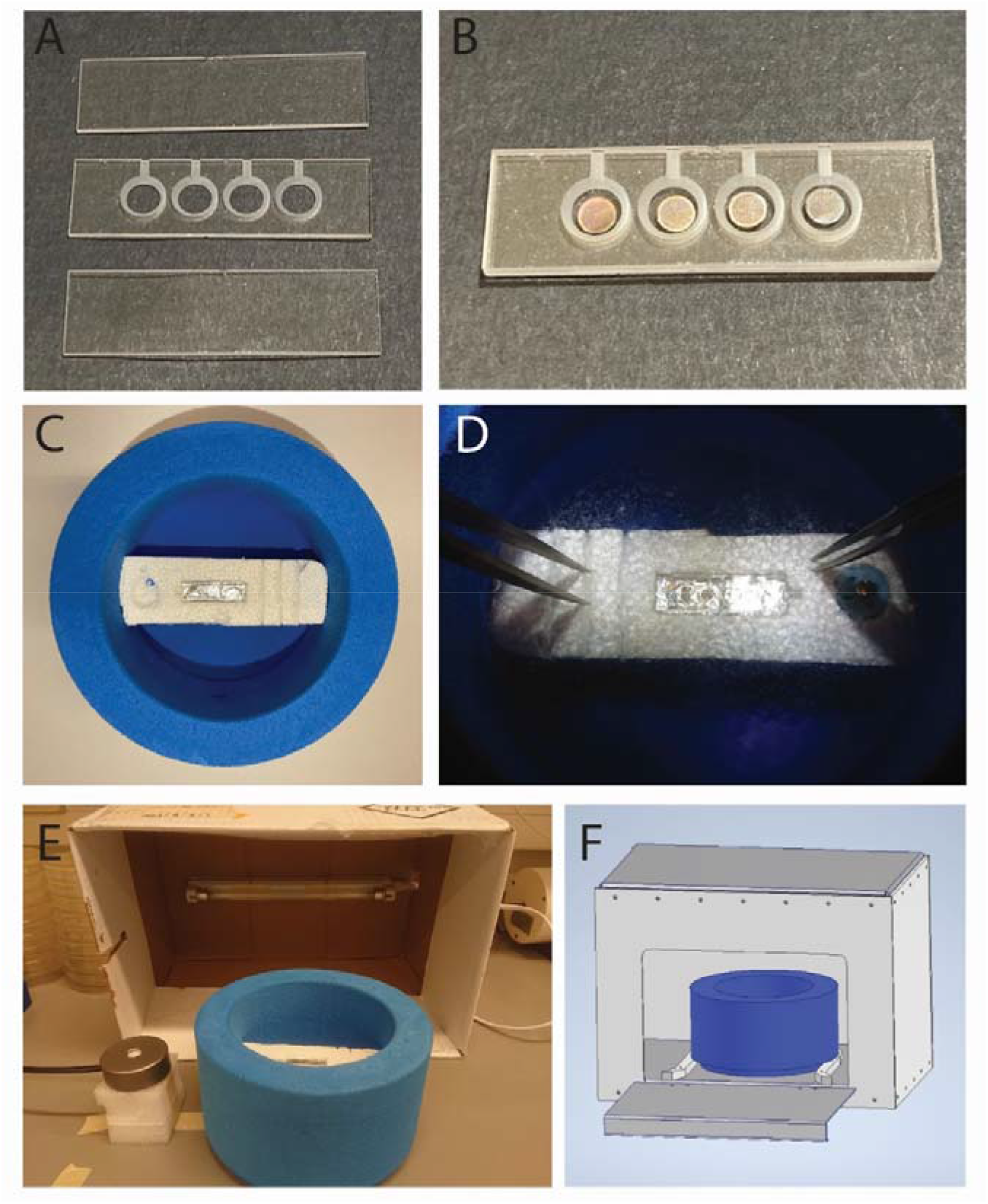
Details of UV inactivation device. Sample containing grids can be loaded into a quartz glass grid holder (unassembled (A) and assembled with grids (B)) under liquid nitrogen conditions. A foam dewar contains a Styrofoam platform for the placement of the quartz glass sample holder during assembly and inactivation, and includes a place for the grid box, an aluminum-lined well for the assemble sample holder (white arrow) and slots for storing each quartz plate (at room temperature (C) and cooled with liquid nitrogen (D)). The initial design of the UV light box mounted the light source to the inside of a cardboard box that could be placed over the cooled sample and UV sensor (E). The final design of the UV light box allows for insertion of the foam dewar into the side (visualized in F).

**Figure S2.**
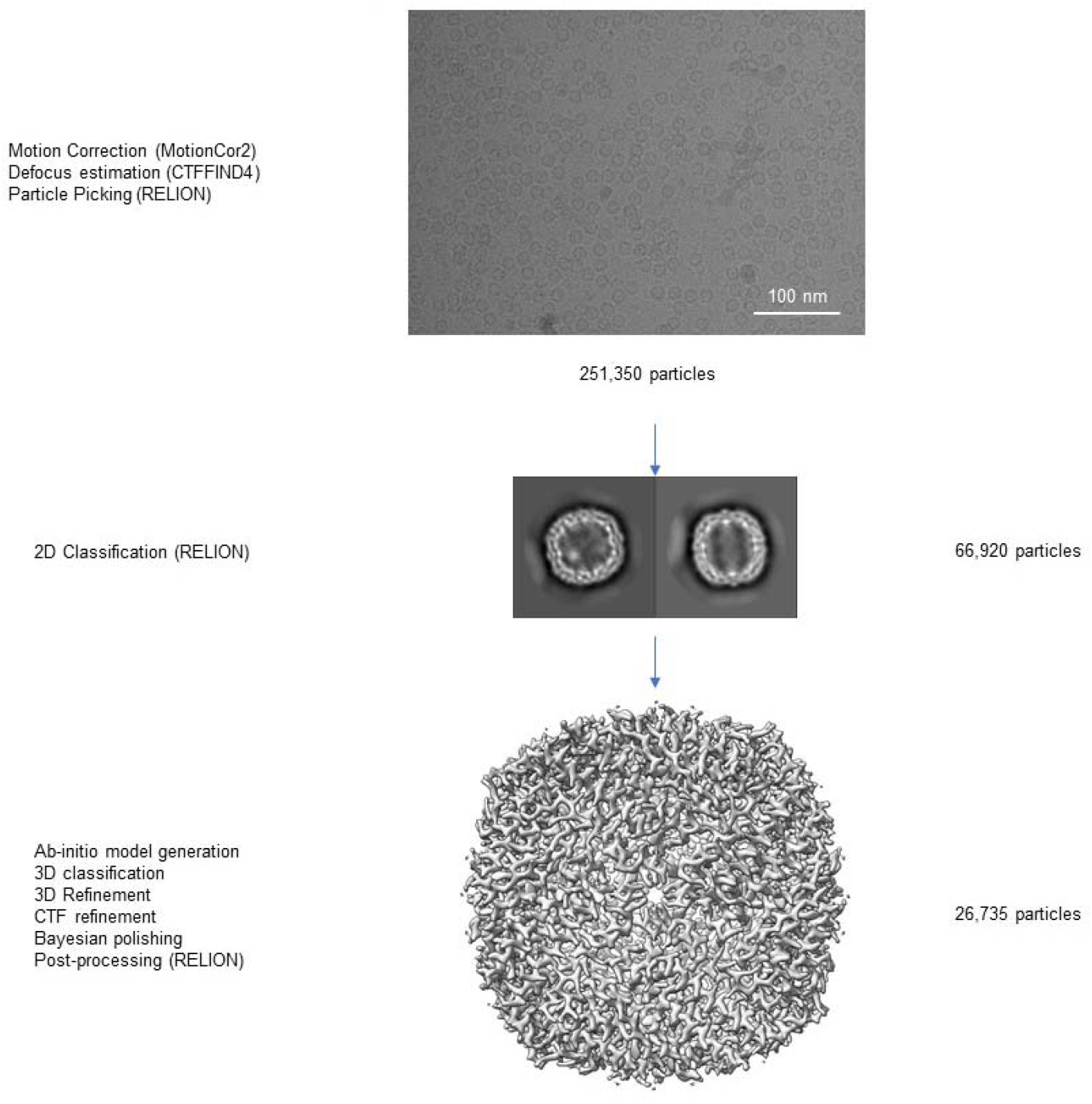
Cryo-EM data processing workflow. A typical micrograph shown as well as representatives 2D classes. 3D classification was performed to distinguish heterogeneity in the sample and the final refined map is showed.

**Figure S3.**
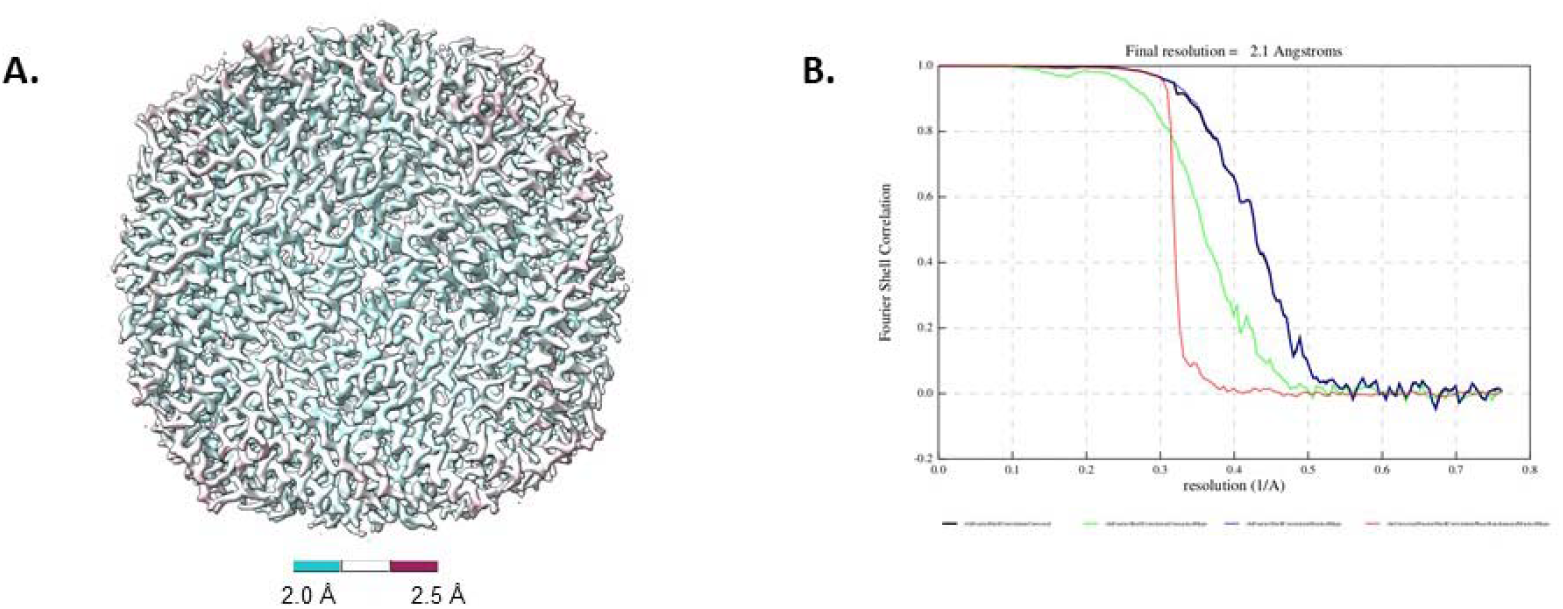
Resolution assessment of cryo-EM structure. A. Local resolution map B. Global resolution assessment by Fourier shell correlation at the 0.143 criterion

**Table S1.** Cryo-EM data collection parameters and processing.

|  | Apoferritin |
| --- | --- |
| **Magnification** | 130,000 |
| **Pixel size (Å)** | 0.328 |
| **Voltage (kV)** | 300 |
| **Electron exposure (e<sup>-</sup>/Å<sup>2</sup>)** | 50 |
| **Frames** | 50 |
| **Defocus Range (μm)** | -0.5 to -2.5 |
| **Number of movies collected** | 1040 |
| **Particles extracted** | 251,350 |
| **Final** | 26,735 |
| **Symmetry** | O |
| **Masked resolution at FSC=0.143 (Å)** | 2.1 |
| **Map sharpening B-factor (Å<sup>2</sup>)** | -44 |

## References

Barth, Z. K., Netter, Z., Angermeyer, A., Bhardwaj, P., & Seed, K. D. (2020). A Family of Viral Satellites Manipulates Invading Virus Gene Expression and Can Affect Cholera Toxin Mobilization. MSystems, 5(5). https://doi.org/10.1128/msystems.00358-20

Berger, C., Ravelli, R. B. G., López-Iglesias, C., Kudryashev, M., Diepold, A., & Peters, P. J. (2021). Structure of the Yersinia injectisome in intracellular host cell phagosomes revealed by cryo FIB electron tomography. Journal of Structural Biology, 213(1). https://doi.org/10.1016/j.jsb.2021.107701

Boyce, J. M., & Donskey, C. J. (2019). Understanding ultraviolet light surface decontamination in hospital rooms: A primer. Infection Control and Hospital Epidemiology, 40(9), 1030–1035. https://doi.org/10.1017/ice.2019.161

Britt, A. B. (1996). DNA damage and repair in plants. Annual Review of Plant Physiology and Plant Molecular Biology, 47(1), 75–100. https://doi.org/10.1146/annurev.arplant.47.1.75

Buonanno, M., Welch, D., Shuryak, I., & Brenner, D. J. (2020). Far-UVC light (222 nm) efficiently and safely inactivates airborne human coronaviruses. Scientific Reports, 10(1), 1–8. https://doi.org/10.1038/s41598-020-67211-2

Caspar, D., & Klug, A. (1962). Physical Principles in the Construction of Regular Viruses. Cold Spring Harb Symp Quant Biol, 27, 1–24. https://doi.org/doi:10.1101/SQB.1962.027.001.005

Castaño-Díez, D., Kudryashev, M., Arheit, M., & Stahlberg, H. (2012). Dynamo: A flexible, userfriendly development tool for subtomogram averaging of cryo-EM data in high-performance computing environments. Journal of Structural Biology, 178(2), 139–151. https://doi.org/10.1016/j.jsb.2011.12.017

Castaño-Díez, D., Kudryashev, M., & Stahlberg, H. (2017). Dynamo Catalogue: Geometrical tools and data management for particle picking in subtomogram averaging of cryo-electron tomograms. Journal of Structural Biology, 197(2), 135–144. https://doi.org/10.1016/j.jsb.2016.06.005

Chen, Z., Sun, L., Zhang, Z., Fokine, A., Padilla-Sanchez, V., Hanein, D., Jiang, W., Rossmann, M. G., & Rao, V. B. (2017). Cryo-EM structure of the bacteriophage T4 isometric head at 3.3-Å resolution and its relevance to the assembly of icosahedral viruses. Proceedings of the National Academy of Sciences of the United States of America, 114(39), E8184–E8193. https://doi.org/10.1073/pnas.1708483114

Diebolder, C. A., Dillard, R. S., & Renault, L. L. R. (2021). From tube to structure: SPA cryo-EM workflow using apoferritin as an example. Methods in Molecular Biology, 2305, 229–256. https://doi.org/10.1007/978-1-0716-1406-8_12

Durchschlag, H., Fochler, C., Feser, B., Hausmann, S., Seroneit, T., Swientek, M., Swoboda, E., Winklmair, A., Wlček, C., & Zipper, P. (1996). Effects of X-and UV-irradiation on proteins. Radiation Physics and Chemistry, 47(3), 501–505. https://doi.org/10.1016/0969-806X(95)00138-N

Ferraro, G., Ciambellotti, S., Messori, L., & Merlino, A. (2017). Cisplatin Binding Sites in Human H-Chain Ferritin. Inorganic Chemistry, 56(15), 9064–9070. https://doi.org/10.1021/acs.inorgchem.7b01072

Goddard, T. D., Huang, C. C., & Ferrin, T. E. (2005). Software extensions to UCSF chimera for interactive visualization of large molecular assemblies. Structure, 13(3), 473–482. https://doi.org/10.1016/j.str.2005.01.006

Goddard, T. D., Huang, C. C., Meng, E. C., Pettersen, E. F., Couch, G. S., Morris, J. H., & Ferrin, T. E. (2018). UCSF ChimeraX: Meeting modern challenges in visualization and analysis. Protein Science, 27(1), 14–25. https://doi.org/10.1002/pro.3235

Hagen, W. J. H., Wan, W., & Briggs, J. A. G. (2017). Implementation of a cryo-electron tomography tilt-scheme optimized for high resolution subtomogram averaging. Journal of Structural Biology, 197(2), 191–198. https://doi.org/10.1016/j.jsb.2016.06.007

Humphrey, W., Dalke, A., & Schulten, K. (1996). VMD: Visual Molecular Dynamics. Journal of Molecular Graphics, 14(October 1995), 33–38. https://www.tapbiosystems.com/tap/products/index.htm

IES. (2020). IES Committee Reports□: Germicidal Ultraviolet (GUV) – Frequently Asked Questions. Illuminating Engineering Society, 1–24. https://media.ies.org/docs/standards/IESCR-2-20-V1a-20200507.pdf%0A https://www.ies.org/standards/committee-reports/.Kevin

Jin, Q., Vogt, S., Lai, B., Chen, S., Finney, L., Gleber, S. C., Ward, J., Deng, J., Mak, R., Moonier, N., & Jacobsen, C. (2015). Ultraviolet germicidal irradiation and its effects on elemental distributions in mouse embryonic fibroblast cells in x-ray fluorescence microanalysis. PLoS ONE, 10(2), 1–14. https://doi.org/10.1371/journal.pone.0117437

Juraszek, J., Rutten, L., Blokland, S., Bouchier, P., Voorzaat, R., Ritschel, T., Bakkers, M. J. G., Renault, L. L. R., & Langedijk, J. P. M. (2021). Stabilizing the closed SARS-CoV-2 spike trimer. Nature Communications, 12(1), 1–8. https://doi.org/10.1038/s41467-020-20321-x

Kremer, J. R., Mastronarde, D. N., & McIntosh, J. R. (1996). Computer visualization of threedimensional image data using IMOD. Journal of Structural Biology, 116(1), 71–76. https://doi.org/10.1006/jsbi.1996.0013

Krisko, A., & Radman, M. (2010). Protein damage and death by radiation in Escherichia coli and Deinococcus radiodurans. Proceedings of the National Academy of Sciences of the United States of America, 107(32), 14373–14377. https://doi.org/10.1073/pnas.1009312107

Kucukelbir, A., Sigworth, F. J., & Tagare, H. D. (2014). Quantifying the local resolution of cryo-EM density maps. Nature Methods, 11(1), 63–65. https://doi.org/10.1038/nmeth.2727

Lorenz, C. M., Wolk, B. M., Quan, C. P., Alcala, E. W., Eng, M., McDonald, D. J., & Matthews, T. C. (2009). The effect of low intensity ultraviolet-c light on monoclonal antibodies. Biotechnology Progress, 25(2), 476–482. https://doi.org/10.1002/btpr.157

Ma, J., Su, D., Sun, Y., Huang, X., Liang, Y., Fang, L., Ma, Y., Li, W., Liang, P., & Zheng, S. (2021). Cryo-EM structure of S-Trimer, a subunit vaccine candidate for COVID-19. Journal of Virology, March. https://doi.org/10.1128/jvi.00194-21

Mastronarde, D. N. (2005). Automated electron microscope tomography using robust prediction of specimen movements. Journal of Structural Biology, 152(1), 36–51. https://doi.org/10.1016/j.jsb.2005.07.007

Mastronarde, D. N., & Held, S. R. (2017). Automated tilt series alignment and tomographic reconstruction in IMOD. Journal of Structural Biology, 197(2), 102–113. https://doi.org/10.1016/j.jsb.2016.07.011

Muok, A. R., Ortega, D. R., Kurniyati, K., Yang, W., Maschmann, Z. A., Sidi Mabrouk, A., Li, C., Crane, B. R., & Briegel, A. (2020). Atypical chemoreceptor arrays accommodate high membrane curvature. Nature Communications, 11(1), 1–13. https://doi.org/10.1038/s41467-020-19628-6

Panikkanvalappil, S. R., Hira, S. M., & El-Sayed, M. A. (2016). Elucidation of ultraviolet radiation-induced cell responses and intracellular biomolecular dynamics in mammalian cells using surface-enhanced Raman spectroscopy. Chemical Science, 7(2), 1133–1141. https://doi.org/10.1039/c5sc03817k

Pattison, D. I., & Davies, M. J. (2006). Actions of ultraviolet light on cellular structures. In L. P. Bignold (Ed.), Cancer: Cell Structures, Carcinogens and Genomic Instability (Issue 96, pp. 131–157). Birkhäuser Verlag/Switzerland. https://doi.org/10.1007/3-7643-7378-4_6

Pettersen, E. F., Goddard, T. D., Huang, C. C., Couch, G. S., Greenblatt, D. M., Meng, E. C., & Ferrin, T. E. (2004). UCSF Chimera - A visualization system for exploratory research and analysis. Journal of Computational Chemistry, 25(13), 1605–1612. https://doi.org/10.1002/jcc.20084

Phillips, J. C., Braun, R., Wang, W., Gumbart, J., Tajkhorshid, E., Villa, E., Chipot, C., Skeel, R. D., Kalé, L., & Schulten, K. (2005). Scalable molecular dynamics with NAMD. Journal of Computational Chemistry, 26(16), 1781–1802. https://doi.org/10.1002/jcc.20289

Rohou, A., & Grigorieff, N. (2015). CTFFIND4: Fast and accurate defocus estimation from electron micrographs. Journal of Structural Biology, 192(2), 216–221. https://doi.org/10.1016/j.jsb.2015.08.008

Scheres, S. H. W. (2012). A bayesian view on cryo-EM structure determination. Journal of Molecular Biology, 415(2), 406–418. https://doi.org/10.1016/j.jmb.2011.11.010

Silva-Valenzuela, C. A., & Camilli, A. (2019). Niche adaptation limits bacteriophage predation of Vibrio cholerae in a nutrient-poor aquatic environment. Proceedings of the National Academy of Sciences, 116(5), 1627–1632. https://doi.org/10.1073/pnas.1810138116

Singharoy, A., Teo, I., McGreevy, R., Stone, J. E., Zhao, J., & Schulten, K. (2016). Molecular dynamics-based refinement and validation for sub-5 Å cryo-electron microscopy maps. ELife, 5(JULY), 1–33. https://doi.org/10.7554/eLife.16105

Subramaniam, S. (2020). COVID-19 and cryo-EM. IUCrJ, 7, 575–576. https://doi.org/10.1107/S2052252520008799

Welch, D., Buonanno, M., Grilj, V., Shuryak, I., Crickmore, C., Bigelow, A. W., Randers-Pehrson, G., Johnson, G. W., & Brenner, D. J. (2018). Far-UVC light: A new tool to control the spread of airborne-mediated microbial diseases. Scientific Reports, 8(1), 1–7. https://doi.org/10.1038/s41598-018-21058-w

Wolff, G., Limpens, R. W. A. L., Zevenhoven-Dobbe, J. C., Laugks, U., Zheng, S., de Jong, A. W. M., Koning, R. I., Agard, D. A., Grünewald, K., Koster, A. J., Snijder, E. J., & Bárcena, M. (2020). A molecular pore spans the double membrane of the coronavirus replication organelle. Science, 369(6509), 1395–1398. https://doi.org/10.1126/SCIENCE.ABD3629

Xiong, Q., Morphew, M. K., Schwartz, C. L., Hoenger, A. H., & Mastronarde, D. N. (2009). CTF determination and correction for low dose tomographic tilt series. Journal of Structural Biology, 168(3), 378–387. https://doi.org/10.1016/j.jsb.2009.08.016

Yang, J., Anishchenko, I., Park, H., Peng, Z., Ovchinnikov, S., & Baker, D. (2020). Improved protein structure prediction using predicted interresidue orientations. Proceedings of the National Academy of Sciences of the United States of America, 117(3), 1496–1503. https://doi.org/10.1073/pnas.1914677117

Yang, W., Keith Cassidy, C., Ames, P., Diebolder, C. A., Schulten, K., Luthey-Schulten, Z., Parkinson, J. S., & Briegel, A. (2019). In situ conformational changes of the escherichia coli serine chemoreceptor in different signaling states. MBio, 10(4).https://doi.org/10.1128/mBio.00973-19

Yao, H., Song, Y., Chen, Y., Wu, N., Xu, J., Sun, C., Zhang, J., Weng, T., Zhang, Z., Wu, Z., Cheng, L., Shi, D., Lu, X., Lei, J., Crispin, M., Shi, Y., Li, L., & Li, S. (2020). Molecular Architecture of the SARS-CoV-2 Virus. Cell, 183(3), 730–738.e13. https://doi.org/10.1016/j.cell.2020.09.018

Zhang, K. (2016). Gctf: Real-time CTF determination and correction. Journal of Structural Biology, 193(1), 1–12. https://doi.org/10.1016/j.jsb.2015.11.003

Zheng, S. Q., Palovcak, E., Armache, J. P., Verba, K. A., Cheng, Y., & Agard, D. A. (2017). MotionCor2: Anisotropic correction of beam-induced motion for improved cryo-electron microscopy. Nature Methods, 14(4), 331–332. https://doi.org/10.1038/nmeth.4193

Zivanov, J., Nakane, T., Forsberg, B. O., Kimanius, D., Hagen, W. J. H., Lindahl, E., & Scheres, S. H. W. (2018). New tools for automated high-resolution cryo-EM structure determination in RELION-3. ELife, 7, 1–22. https://doi.org/10.7554/eLife.42166

Zivanov, J., Nakane, T., & Scheres, S. H. W. (2019). A Bayesian approach to beam-induced motion correction in cryo-EM single-particle analysis. IUCrJ, 6, 5–17. https://doi.org/10.1107/S205225251801463X

Zivanov, J., Nakane, T., & Scheres, S. H. W. (2020). Estimation of high-order aberrations and anisotropic magnification from cryo-EM data sets in RELION-3.1. IUCrJ, 7, 253–267. https://doi.org/10.1107/S2052252520000081

